# Genetic analyses reveal distinct functional roles of Eut microcompartment shell proteins in ethanolamine metabolism in *Salmonella*

**DOI:** 10.64898/2026.10.07.757396

**Authors:** Minal R. Bhalerao, Ajit A. Sutar, Aishwarya S. Davkhar, Yashodhara D. Shinde, Sachin B. Agawane, Chiranjit Chowdhury

## Abstract

Ethanolamine utilization is an important metabolic capability of enteric bacteria and has been associated with bacterial fitness and adaptation within diverse host environment. In *Salmonella enterica* serovar Typhimurium, ethanolamine is metabolized within the bacterial microcompartment (Eut MCP), which is composed of five shell protein subunits. Deciphering how self-assembled shell proteins spatially organize the pathway, retain metabolic intermediates and create proper environment for catalysis is crucial for understanding role of Eut MCP in cellular metabolism and pathogenesis. Through genetic mutant construction, growth and metabolic assays, microscopy, and metabolite quantification, we demonstrate that each shell protein fulfills a distinct role to the functioning of the Eut MCP. Our findings provide mechanistic insights into the structural organization and functional dynamics of the bacterial microcompartment organelle.

## 1. Introduction

Bacterial microcompartments (MCPs) are sophisticated protein-based organelles that encapsulate enzymes involved in specific metabolic pathways ranging from carbon dioxide fixation, plant sugar degradation, aromatic degradation and the metabolism linked to several diseases, including cancer and heart disease (Bhalerao *et al*., 2025, Chowdhury *et al*., 2014, Doron *et al*., 2023, Herring *et al*., 2018, Kerfeld & Erbilgin, 2015, Prentice, 2021, Stewart *et al*., 2021). The protein shells of MCPs are polyhedral in structure that ranges from 80 to 200 nm in cross-section (Chowdhury *et al*., 2014, Yeates *et al*., 2013), comprising a conserved set of shell proteins of 5 to 10 different types, unlike other biological condensates that assemble from a single type of shell protein (Cheng *et al*., 2011, Gupta & Uversky, 2026, Timane & Chowdhury, 2026). The shell proteins of bacterial MCP mainly belong to three protein family types that primarily form facets and vertices of the polyhedral MCP shell (Cheng *et al*., 2011, Ochoa & Yeates, 2021, Yeates *et al*., 2011). The first type comprises single BMC domain (Pfam00936) that assemble into a cyclic hexamer. The shell protein consisting tandem BMC proteins (fusion of two Pfam00936 domains) represents the second type, which organizes into pseudohexameric trimer. The third one is BMV family (Pfam PF03319), a non BMC homolog that forms pentamer (Chowdhury *et al*., 2014, Stewart *et al*., 2021). Some MCP shell proteins also harbor unique features that contain either N-terminal beta-barrel, C-terminal appendages or an iron-sulfur cluster down the center of hexamer, in addition to their canonical conformation (Cheng *et al*., 2011, Chowdhury *et al*., 2014, Crowley *et al*., 2008, Tanaka *et al*., 2010). In some MCPs there also exists unique shell proteins that lack recognizable BMC, or BMV domains (Cheng *et al*., 2011, Sinha *et al*., 2012). Each of these diverse shell protein components are suggested to serve unique functional attributes that maintain proper size, shape of MCPs and spatial organization to create optimal conditions for metabolic functions (Ke *et al*., 2026, Timane & Chowdhury, 2026, Yang *et al*., 2025). Some shell proteins are actively involved in selectively controlling the movement of substrates, cofactors and products, restrict outward diffusion of volatile and/or toxic intermediates thus preventing cellular toxicity and/or reduce carbon loss (Cheng *et al*., 2011, Chowdhury *et al*., 2015, Chowdhury *et al*., 2014, Crowley *et al*., 2010, Hartzler *et al*., 2026, Ochoa *et al*., 2023, Tanaka *et al*., 2010), or implicate in recruiting enzyme cargo within MCP lumen (Choudhary *et al*., 2012, Fan & Bobik, 2011, Fan *et al*., 2012, Held *et al*., 2016, Lehman *et al*., 2017, Yang *et al*., 2025). Nevertheless, there are still many unanswered questions on individual roles of shell proteins in MCP biogenesis and metabolism, albeit availability of gross structural insights (Chowdhury *et al*., 2015, Chowdhury *et al*., 2016, Crowley *et al*., 2010, Crowley *et al*., 2008, Hartzler *et al*., 2026, Stewart *et al*., 2021, Tanaka *et al*., 2010, Yeates *et al*., 2013).

*Salmonella enterica* serovar Typhimurium LT2 (henceforth denoted as *Salmonella*) deploys two types of MCPs to metabolize 1, 2-propanediol and ethanolamine (EA) inside the human gut. Metabolism within MCP promotes pathogen adaptation in host environments and alters gut microbial ecology (Bhalerao *et al*., 2025, Prentice, 2021, Stewart *et al*., 2020, Corcoran & Skaar, 2026). In recent past, the MCP shells of both the types have been exploited to serve as sub-cellular nanoreactor in several biotechnology applications, where synthetic MCP shell has been constructed with minimal number of important shell proteins (Held *et al*., 2016, Held *et al*., 2013, Lawrence *et al*., 2014, Lee *et al*., 2019, Parsons *et al*., 2010, Pokhrel *et al*., 2021, Shinde & Chowdhury, 2024). Therefore, considerable attention has been directed toward elucidating the native mechanisms underlying MCP assembly and function.

Earlier, genetic study suggested role of individual shell proteins of propanediol utilization microcompartment (Pdu MCP) in maintaining polyhedral morphology and MCP metabolism (Cheng *et al*., 2011, Chowdhury *et al*., 2016, Kennedy *et al*., 2022, Lehman *et al*., 2017). Later, Pdu MCPs were purified and significance of individual shell proteins were thoroughly evaluated (Chowdhury *et al*., 2015, Havemann & Bobik, 2003, Kennedy *et al*., 2021, Kennedy *et al*., 2022, Sinha *et al*., 2012, Yang *et al*., 2022). Unlike Pdu system, the specific roles of shell proteins of Eut MCP are uncertain, and the effects of deleting these genes singly on ethanolamine degradation are also elusive. *Salmonella* Eut MCPs have not yet been purified, hindering quantitative assessment of shell component abundance and their functional contributions to MCP assembly and ethanolamine metabolism. In *Salmonella*, the shell of Eut MCP is composed of five shell proteins, namely EutS, M, N, L, K, encoded in a 17-gene operon known as the ethanolamine utilization operon (*eut*) (Fig. 1) (Kofoid *et al*., 1999). In the functional model, ethanolamine enters the lumen of MCPs after crossing the outer protein shell (Fig. 1). The ethanolamine is converted to acetaldehyde intermediate within the lumen by a coenzyme B_12_ dependent enzyme, ethanolamine ammonia lyase (EutBC). Acetaldehyde further reacts with aldehyde dehydrogenase (EutE) to produce acetyl-CoA, which later joins TCA cycle in the central metabolism (Chowdhury *et al*., 2014, Garsin, 2010). Prior studies with *Salmonella* showed that a quadruple mutant lacking EutMNLK was failed to grow on EA containing media (Brinsmade *et al*., 2005) and the impaired growth might be attributed to poor retention of the volatile acetaldehyde intermediate within the mutant MCP shell (Penrod & Roth, 2006). Recently, mutants lacking single shell protein exhibited severe growth defect (Yang *et al*., 2025) and there were spatial separation of shell and cargo enzymes in some of the shell protein mutants (Yang *et al*., 2025, Held *et al*., 2016).

**Figure 1.**
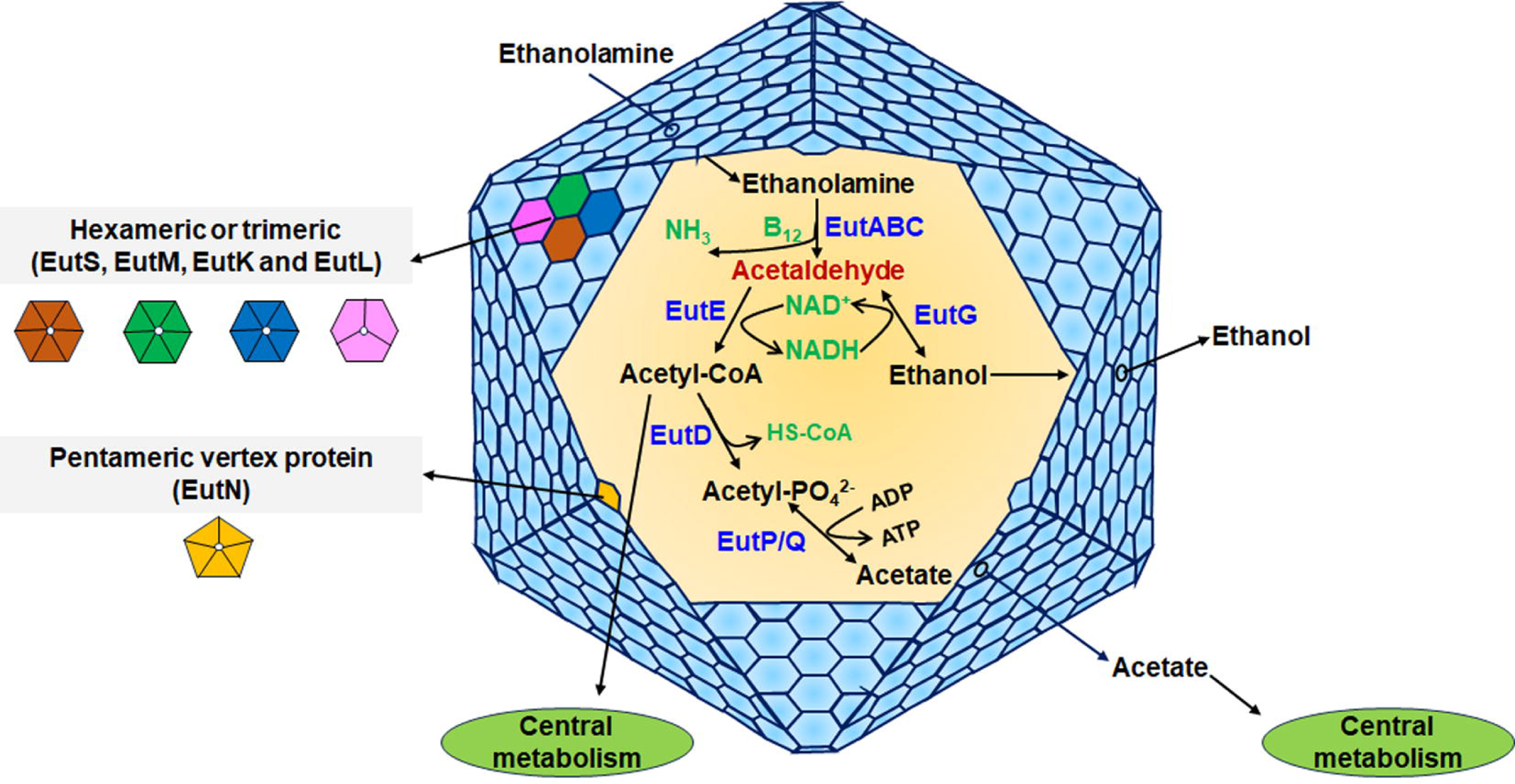
Schematic model of the Eut MCP and EA utilization pathway.

To better understand the specific functional roles of the individual shell proteins of Eut MCP, we constructed in-frame deletion mutants of each *eut* shell protein gene and determined their effects on MCP morphology and ethanolamine catabolism using microscopic analyses, growth studies, estimation of metabolites and enzyme activity in crude cell lysate. Current study would provide insights into operational principle of Eut MCP shell proteins in EA metabolism in *Salmonella*.

## Results

### Mutation in shell protein results aberrant MCP structure

Both transmission electron microscopy (TEM) of thin-sectioned bacterial sample and confocal microscopy were used to examine the effects of the individual shell protein deletion mutations on the structure of the Eut MCP of *Salmonella*.

In the absence of EA and B_12_, no endogenous Eut proteins were expressed in *Salmonella* and therefore, there were no polyhedral structure were seen (Fig. 2a). MCPs in *eutK*, -*M* and -*L* deletion mutants resulted in the formation of what appeared to be polar bodies. As compared to *eutM*, the polar aggregates in *eutL* and *eutK* mutants were smaller in size (Fig. 2). In some *eutK* mutant, some protein bodies were appeared at both side of the pole (Fig. 2). Unlike WT and other mutants, an *eutN* mutant generated MCP assemblies that were elongated and spanned within two cells (Fig. 2). An *eutS* deletion had no obvious effects on MCP formation and the mutant cells mostly formed MCPs similar to wild type (Fig. 2).

**Figure 2.**
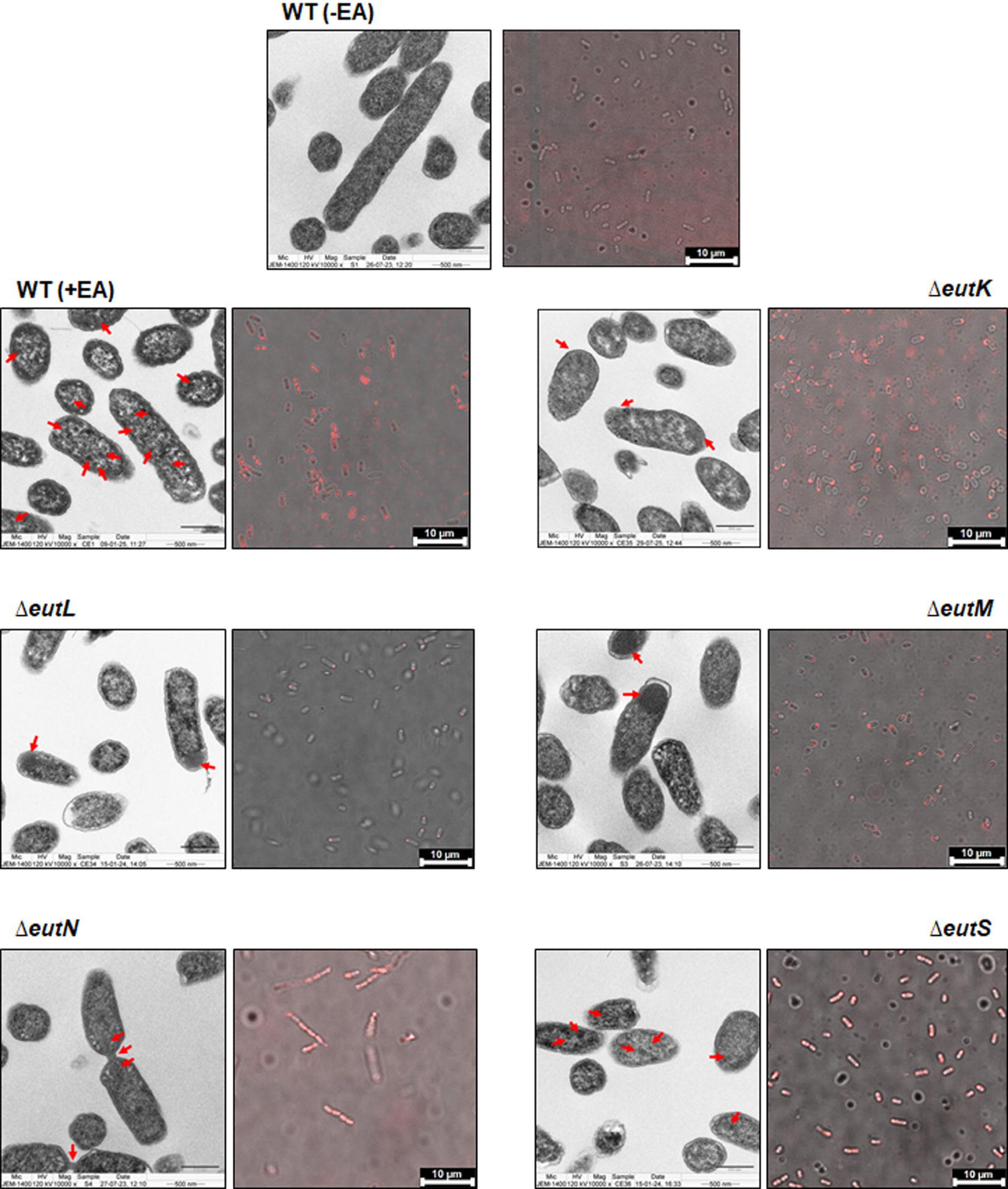
Transmission electron microscopy (TEM) of ultra-thin section and confocal microscopy of the wild type *S*. Typhimurium and Eut shell protein mutants grown in the presence of EA and B_12_, Scale bar: 500 nm and 10 µm. TEM determines morphological aberration of the MCP upon genetic mutation. The confocal microscopy determines localization and distribution of the enzyme (EutBC) tagged with m-Cherry reporter.

While TEM analyses demonstrated morphological insights of both WT and shell protein mutants, the confocal laser scanning microscopy has been carried out next to track protein localization and dynamics.

Since EutBC is a localized within Eut MCP (Costa & Escalante-Semerena, 2022), we reasoned that spatial organization could be better visualized by tagging EutBC with a reporter protein chromosomally. In this context, an m-Cherry reporter protein has been fused at R298 of EutC in the chromosomal loci (details in experimental procedure below). The WT cells grown in the glycerol media without EA and B_12_ did not produce any fluorescent signal, as expected (Fig. 2a). However, in presence of EA and B_12_, fluorescent punctate signal was emerged throughout the cytoplasm. An *eutM* mutant generated large bright puncta at the one side of the pole, similar to large protein aggregate as observed in TEM image. In contrast, the polar puncta were smaller and less bright in case of *eutK* and *eutL* mutant (Fig. 2). In some *eutK* mutant cells, the fluorescent dots were observed at both the side of the pole (Fig. S1). In case of *eutS* mutant, punctate fluorescence were distributed throughout the cell, almost similar to that of wild-type (Fig. 2, Fig. S1). An *eutN* mutant, in contrast produced elongated cell. There were no puncta in *eutN* mutant cell, rather it was what appeared to be a fluorescent rod spanned within multiple bacterial cells (Fig. 2), which was consistently observed in transmission electron microscopic images of the thin sectioned bacterial cell (Fig. 2).

### Growth is impaired in shell protein deletion mutants

To decipher if the deformed shells resulting from shell protein mutation affect EA utilization, the growth of the bacteria was monitored over a period of time in microplate spectrophotometer. To examine if the shell protein mutants are defective EA permeability, the growth assay was performed in no carbon-E (NCE) with EA (30 mM) as a carbon source and was compared at both limiting level (20 nM) and saturating level (150 nM) of vitamin B_12_ (Chowdhury *et al*., 2015, Yang *et al*., 2022, Chowdhury *et al*., 2016). An earlier study of Pdu MCPs showed that substrate permeability defects manifested as growth impairment under limiting B_12_, but these defects were rescued when B_12_ was saturating. During growth at a saturating level of B_12_, the B_12_ dependent dioldehydratase activity become maximized and the MCP shell no longer acts as a physical barrier (Chowdhury *et al*., 2015, Yang *et al*., 2022, Chowdhury *et al*., 2016). Unlike Pdu MCP, the growth of Eut MCP shell protein mutants remained slower in both limiting B_12_ and in saturating B_12_ (Fig. 3a, Fig. S2), indicating potential problem in downstream part of EA metabolism, not in EA uptake. Notably, all shell protein mutants had shown defective growth phenotype (Fig. 3a, Table 1). An *eutM* mutant exhibited slowest growth rate among all mutants (Table 1). This was followed by *eutK, -N, -L* and *eutS* (Fig. 3a, Table 1). In addition, there also found extended lag phase in the growth of all mutants (Fig. 3a). The growth rate of *eutL* was seemed to have corrected after 12 h and grew almost similar fashion to WT (doubling time of WT is 2.93 h, doubling time of *eutL* is 2.99 h) and sometime crossed beyond WT growth. However, growth of the *eutM* and *-K* mutants remained slower throughout the study period (Fig. 3). The slower growth could be attributed to metabolic defect arose from misshapen shell in the mutants. On the other hand, the extended lag phase in bacterial growth might be ascribed to cellular adaptation and repairing process against toxic aldehyde liberating from disrupted shell.

**Figure 3.**
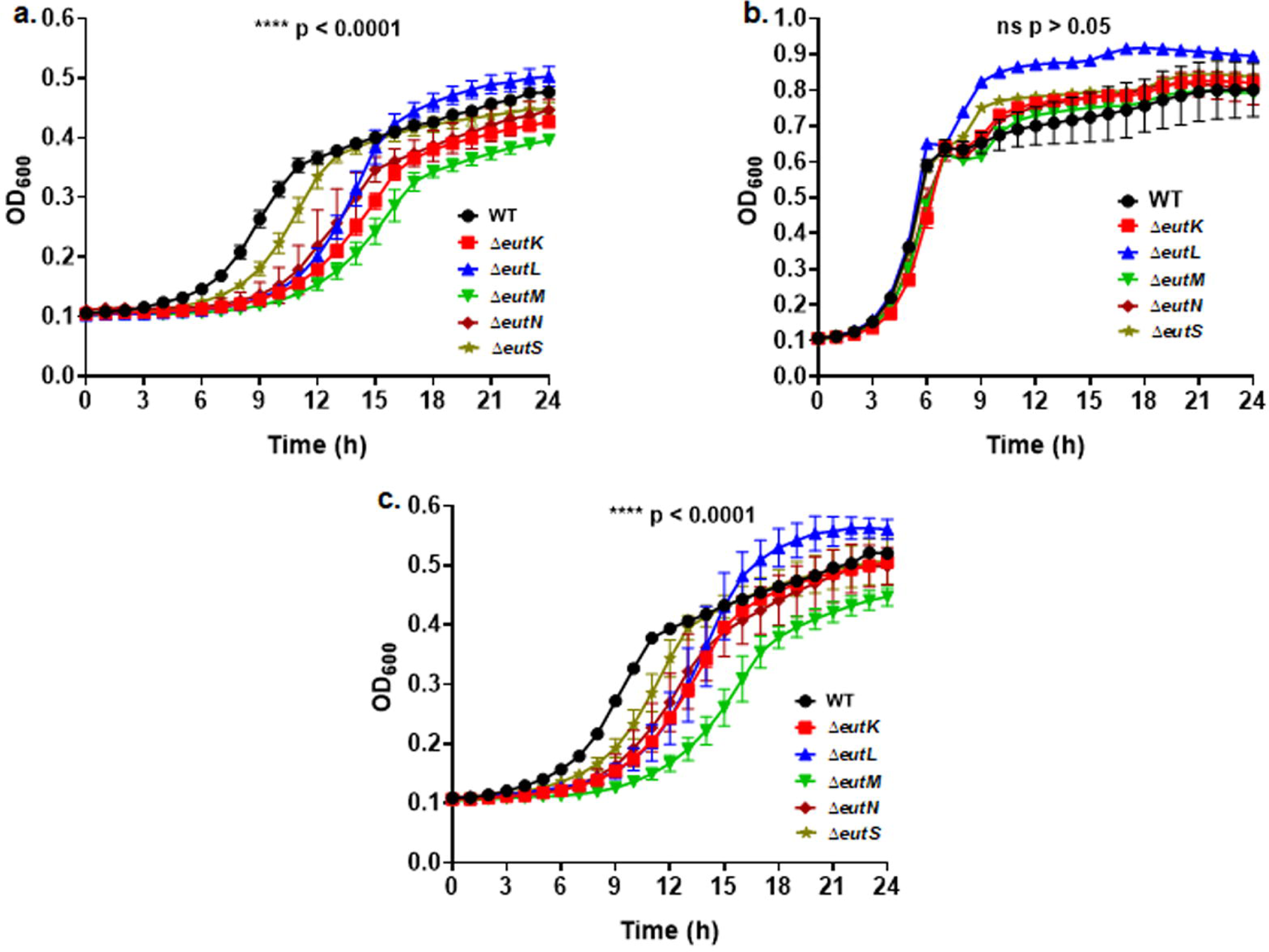
Growth of wild type and Eut shell protein mutants in minimal medium supplemented a) EA as carbon b) EA as nitrogen c) EA as carbon and nitrogen both. Error bars represent ± one standard deviation and are based on three biological replicates. A two-way ANOVA was used for statistical significance ***p < 0.0005, ****p < 0.0001.

**Table 1.** Doubling time of wild type and Eut shell protein mutants in minimal media where EA served as carbon or nitrogen or carbon and nitrogen both.

| Doubling time (h) |  |  |  |
| --- | --- | --- | --- |
| Strain | EA as C | EA as N | EA as C & N |
| WT | 2.93±0.07* | 1.41±0.09<sup>†</sup> | 3.78±0.08 |
| *ΔeutK* | 4.13±0.26 | 1.58±0.19 | 4.19±0.08 |
| *ΔeutL* | 2.99±0.15 | 1.24±0.01 | 3.17±0.18 |
| *ΔeutM* | 4.19±0.05 | 1.48±0.02 | 4.11±0.17 |
| *ΔeutN* | 3.38±0.75 | 1.44±0.01 | 3.99±0.21 |
| *ΔeutS* | 3.09±0.04 | 1.32±0.02 | 3.44±0.18 |

\*Doubling times were calculated from at least three biological replicates measured in triplicate. The error estimate shown is ±one standard deviation. <sup>†</sup>The growth of shell protein mutants were not significantly different from the WT when EA was served as nitrogen source.

*Salmonella* has been known to utilize EA as carbon and nitrogen *in vivo* (Penrod & Roth, 2006, Roof & Roth, 1988, Stojiljkovic *et al*., 1995, Thiennimitr *et al*., 2011). Hence, it would seem reasonable to determine the importance of MCP shell in EA metabolism at different conditions. Both WT and the individual shell protein mutants were grown in no carbon-no nitrogen (NCN) medium, where EA (30 mM) was used as a nitrogen source with glycerol (20 mM), or as a carbon and nitrogen source in NCN medium (Bhalerao *et al*., 2025, Sheppard *et al*., 2004).

Intriguingly, the shell protein mutation had little or no effect when EA was served as nitrogen-source (Fig. 3b); unlike enterohemorrhagic *Escherichia coli*, which utilize EA as nitrogen source (Kendall *et al*., 2012). The growth pattern of shell protein mutants with ethanolamine as both carbon and nitrogen source resembled that observed when ethanolamine served solely as the carbon source (Fig. 3c).

### Eut shell protein mutants produce lower acetate and ethanol

Since, Eut shell protein mutants exhibited defective growth phenotype when EA serves as a carbon source, we presumed potential metabolic deficiencies in carbon utilization. Therefore, we sought to determine acetate and ethanol produced throughout bacterial growth by quantifying their amount accumulated in the media. Previous study suggested that bacteria grown at saturating B_12_ provides a means to measure the extent of aldehyde leakage and product release from the MCP (Chowdhury *et al*., 2015). Acetate and ethanol are key end products of ethanolamine metabolism (Fig. 1), released into the surrounding medium. For experiments where both growth and metabolites were measured, cells were grown in Erlenmeyer flasks to facilitate sampling rather than in a microplate reader as was done in the studies described above (Fig. S3a).

It was observed that *eutK*, *eutS* and *eutL* mutants liberated maximum acetate almost 50% lower as compared to WT (Fig. 4a). The maximum amount of acetate in the media of *eutN* and *eutM* mutants were almost 60% and 30%, respectively to WT (Fig. 4a). For ethanol, maximum levels reached in cultures in *eutS* and *eutM* mutants were almost 50% to that of WT (Fig. 4b). In contrast, *eutK* and -*L* mutants had maximum ethanol only 20% lower than WT; whereas, the amount of ethanol in an *eutN* mutant was almost similar (90%) to WT (Fig. 4b).

**Figure 4.**
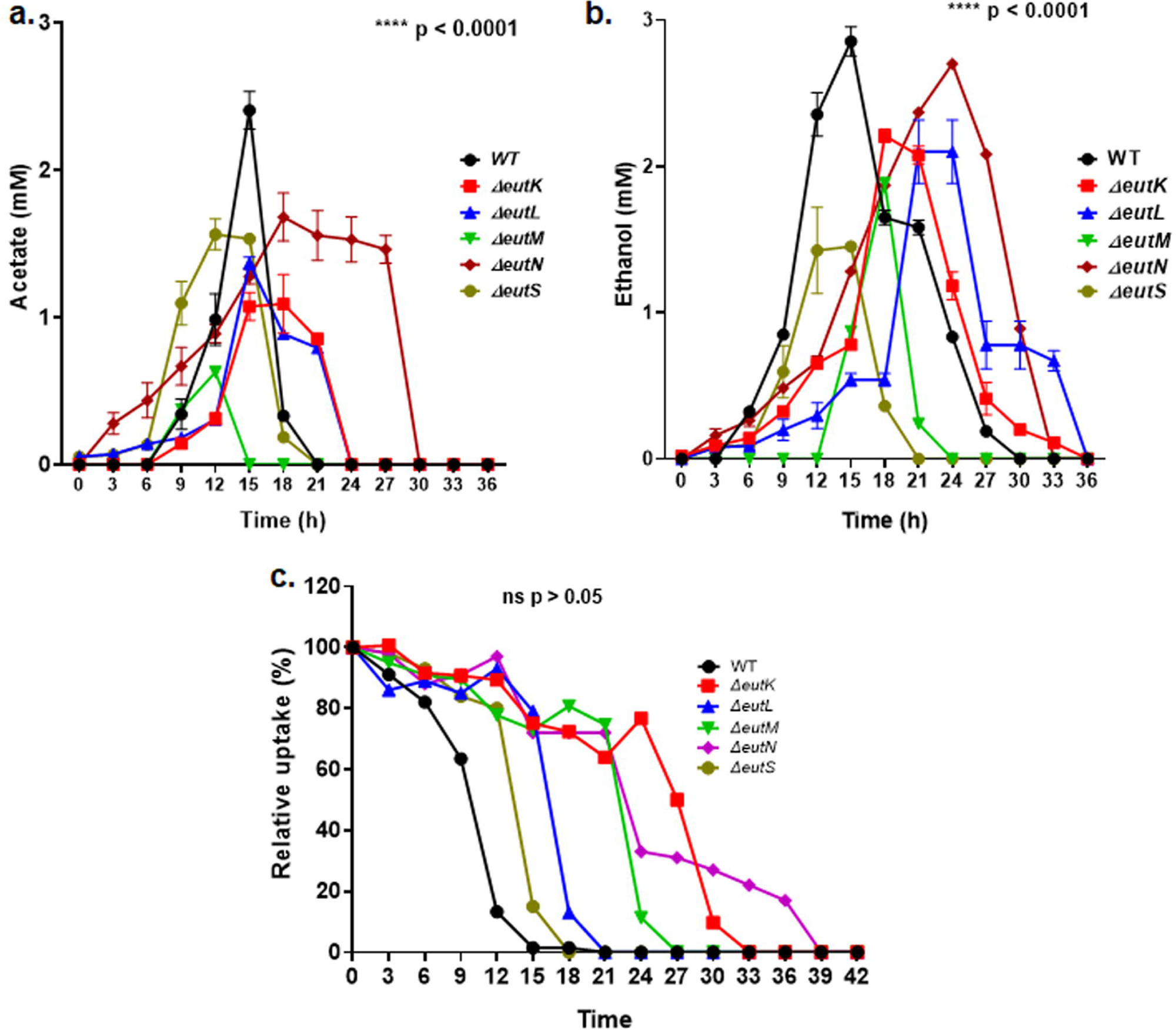
Quantification of products released by wild type *S*. Typhimurium and Eut shell protein mutants grown in presence of EA and B_12_ a) Acetate b) Ethanol. The error bars represent ± one standard deviation and are based on three biological replicates. A two-way ANOVA was used for statistical significance. ****p < 0.0001 compared to the wild type control. c) Quantification of ethanolamine uptake by wild type *S*. Typhimurium and Eut shell protein deletion mutants grown in presence of EA and B_12_. The error bars represent ± one standard deviation and are based on two biological replicates. An unpaired t-test (two-tailed) was used for statistical significance. ns p > 0.05 compared to the wild type control.

Also of note was the finding that all mutants exhibited a delayed onset of acetate and ethanol production, attributable to the lag in their growth initiation.

Here, our analysis of metabolite levels indicated that the level of acetate and ethanol were depleted after reaching to the maximum level (Fig. 4); however, growth continued. Earlier studies depicted that acetate can be assimilated by housekeeping Acs (encoding Acetyl-CoA synthetase) or by combined action of AckA (Acetate kinase) and Pta (Phosphotransacetylase). The ethanol is getting consumed by cytoplasmic AdhP (alcohol dehydrogenase) and Aldh (Aldehyde dehydrogenase) (Huseby & Roth, 2013). Notably, there were rapid consumption of ethanol and acetate by *eutN* mutant (Fig. 4).

### Ethanolamine uptake is normal in the Eut shell protein mutants

To assess if aberrant Eut MCP impeded EA transport through the shell the EA consumption was measured over a period of time.

All individual shell protein deletion mutants displayed EA uptake patterns comparable to the wild type, except the *eutN* mutant (Fig. 4c). We opined that abolishing Eut shell protein had no effect on EA transport. In *eutN* mutant, MCPs conformed rod shaped assembly that were spanned between two or more bacterial cell (Fig. 2). Recent evidence linking membrane morphology to diffusion dynamics suggests that molecules permeate more slowly through rod-shaped structures than through spherical ones of equal volume (Rojas Molina *et al*., 2021), providing a rationale for the reduced EA uptake observed in the *eutN* mutant shell. Earlier, slower substrate uptake was observed in *pduN* mutant as well (Mills *et al*., 2022). Incidentally, intact MCP shell is either icosahedral or spherical in shape (Doron & Kerfeld, 2024, Sutter *et al*., 2019, Yeates *et al*., 2010). As usual the shell protein mutants showed a delay in initiating substrate uptake, consistent with their growth behavior (Fig. 4c).

### Mutant MCP shells are defective in retaining acetaldehyde intermediate within lumen

Inside MCP lumen EA gets converted to acetaldehyde, which should remain inside for the accelerating downstream enzymatic pathways. Our study revealed that Eut shell protein mutants released more acetaldehyde in the media as compared to WT (Fig. 5). For *eutM*, the maximum acetaldehyde level reached to 1.8 mM; whereas in *eutL* the same was 0.7 mM. The *eutS* and *eutK* mutants released considerably less acetaldehyde reaching a maximum of approximately 0.5 mM. Interestingly, *eutN* mutant exhibited a lower peak concentration of acetaldehyde than the other shell protein mutant strains (Fig. 5). The WT MCP had effectively retained the acetaldehyde and the maximum amount found in media was approximately 0.2 mM.

**Figure 5.**
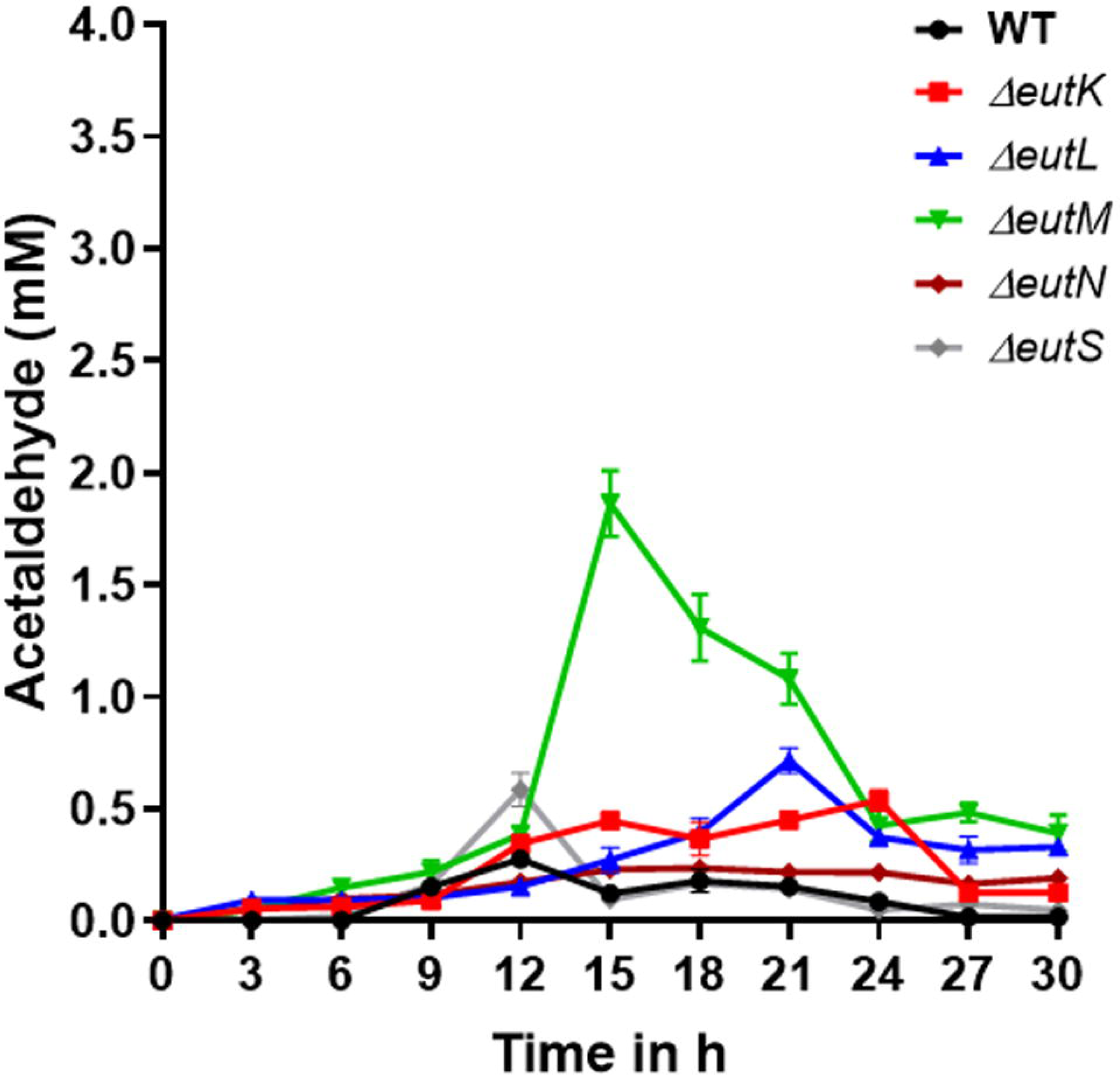
Quantification of acetaldehyde release by wild type *S*. Typhimurium and Eut shell protein mutants grown in presence of EA and B_12_. The error bars represent ± one standard deviation and are based on three biological replicates. A two-way ANOVA was used for statistical significance. **p < 0.005, compared to the wild type control.

A second test was used to better assess the diffusion of acetaldehyde across mutant MCP shell. In the current study, the individual shell protein mutants were grown under a condition where growth was limited by diffusion of acetaldehyde, produced by oxidation of ethanol in the cytoplasm, into the Eut MCP. Prior studies showed that an *eutB* mutant impedes ethanolamine utilization, without obstructing MCP expression (Bhalerao *et al*., 2025); hence, we reasoned that the effects of the shell protein mutation on acetaldehyde diffusion might be more readily observed in an *eutB* background. The current observations support a model in which growth of *Salmonella* on ethanol requires conversion ethanol to acetaldehyde in the cytoplasm by AdhP followed by diffusion of the latter across the MCP shell and into the lumen where it gets converted to acetyl-CoA by the EutE (Chowdhury *et al*., 2015, Huseby & Roth, 2013).

These tests indicated that the relative rate of acetaldehyde movement into the Eut MCP was *eutK* > *eutL* > *eutS* > *eutM* ∼ *eutN* > WT (Fig. S3b).

Remarkably, the rate of aldehyde diffusion varied across five mutant shells when assessed by different methods. In the first case, the acetaldehyde egressed out to the media was measured through direct quantification method. In the second test the same was evaluated on the basis of growth, which was limited by inward diffusion of acetaldehyde and further downstream metabolism.

Given slight variation in the rate of acetaldehyde diffusion through shell protein mutants as estimated by two different methods, both the methods however indicated release of excess acetaldehyde to the media upon deletion of shell proteins.

The high levels of aldehyde released by the shell protein deletion mutants are likely due to an improperly formed MCP that could not retain this small molecule within its lumen. Excessive acetaldehyde release out of the MCP shell might reduce availability carbon intermediate for downstream enzymatic pathway within the MCP lumen, resulting lowering of end product (ethanol and acetate) formation and impeded the growth. Prior studies also showed that a triple shell protein deletion mutant in Cut MCP results in release of higher amounts of acetaldehyde in the media which impairs cell growth (Ochoa *et al*., 2023).

### Enzyme assays confirmed the leaking of key enzymes from the mutant shell

We sought to investigate how deletion of shell protein affects assembly of enzyme within MCP lumen that affects growth and metabolism further. Both *in vivo* model and interactome study proposed interactions between Eut shell protein and lumen enzyme (Held *et al*., 2016, Jallet *et al*., 2024, Jorda *et al*., 2015, Yang *et al*., 2025). A crude enzyme assay from cell lysate indicated that both EAL (EutBC) and aldehyde dehydrogenase (EutE) activities were reduced in all the shell protein deletion mutants, except in *eutS* (Fig. 6). There were more than 80% reduction in EAL activity in the cell devoid of EutK, -M and -N as compared to WT; whereas, the same in *eutL* deletion mutant it was 60% (Fig. 6a). The deletion of *eutL* and *eutM* resulted 40% acetaldehyde dehydrogenase (EutE) activity remaining; whereas, the loss was 40% in case of *eutK* and *eutN* mutants (Fig. 6b). It can be suggested that deletion of shell protein reduced the enzyme loading into MCPs, which often deposited as polar aggregate as observed under microscopic analyses. Unencapsulated cytoplasmic protein aggregates were likely to become solubilized and diluted following cell lysis, thereby diminishing enzymatic activity. Given an *eutS* mutant and WT cells had proper shell closure, the apparent enzyme concentration to the substrate might be higher resulting in higher specific enzyme activity. The higher effective enzyme activity due to crowding effect of the encased enzyme has been depicted elsewhere (Chakraborti *et al*., 2019, Chakraborti *et al*., 2020).

**Figure 6.**
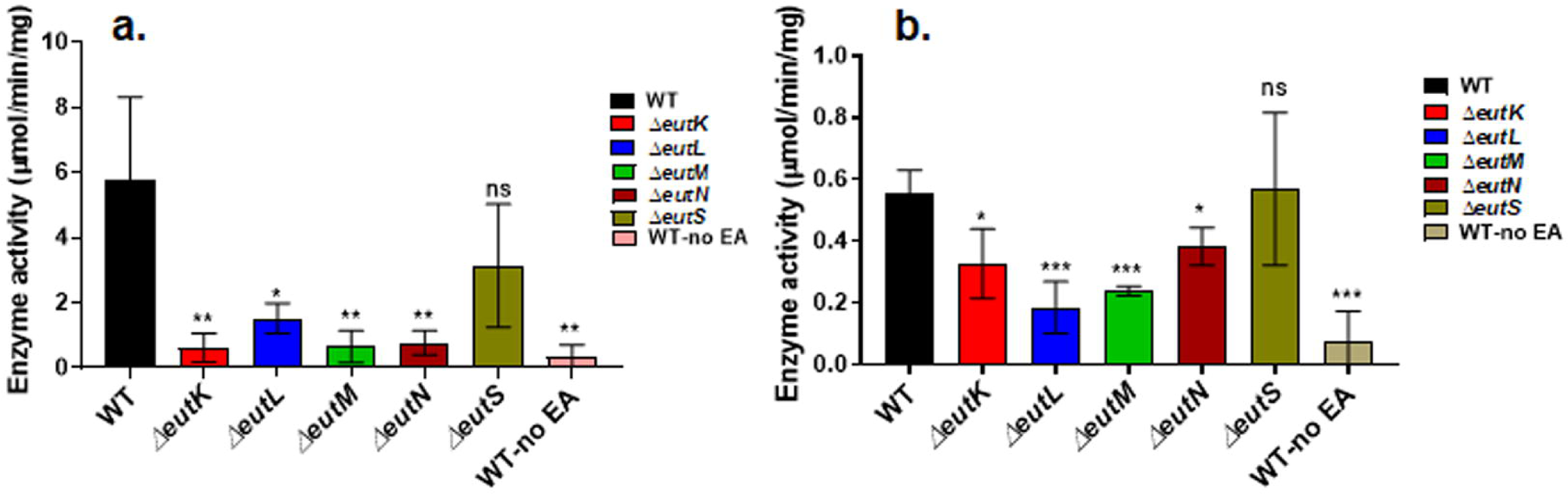
Determination of activity of enzymes from the cell lysate. a) Ethanolamine ammonia lyase (EutBC) b) Acetaldehyde dehydrogenase (EutE) activity from the crude cell lysate. The error bars represent standard deviations. An unpaired t-test (two-tailed) was used for statistical significance as compared to WT. ns p > 0.5, *p < 0.02, **p < 0.05, ***p < 0.005

## Discussion

MCPs are unique proteinaceous organelles that support bacterial survival under distinct nutritional niche. Understanding their biogenesis and function are of considerable interest, to serve both as prospective template for designing novel drug against enteric pathogens and as platforms to get over metabolic bottleneck in biomanufacturing system (Bhalerao *et al*., 2025, Kennedy *et al*., 2022, Lee *et al*., 2019, Prentice, 2021, Jakobson & Tullman-Ercek, 2016). Yet, the native mechanisms governing their assembly and activity remain obscure. Role of shell proteins in MCP metabolism was thoroughly studied in case of Pdu MCP (Chowdhury *et al*., 2015, Chowdhury *et al*., 2016, Fan *et al*., 2012, Havemann *et al*., 2002, Kennedy *et al*., 2021, Kennedy *et al*., 2022, Lehman *et al*., 2017, Sampson & Bobik, 2008, Sinha *et al*., 2012, Yang *et al*., 2022, Mills *et al*., 2022). In contrast, the same in Eut MCP is meager. Here, we systematically generated a series of mutants that lacked individual shell proteins and took deep dive to demonstrate their effect on shell morphology, growth and metabolite time course trajectories *in vivo* followed by *in vitro* enzyme activity.

Deletion of some Eut shell proteins exhibited distinct phenotype than one observed in Pdu system. Microscopic analyses of MCP shell morphology and spatial distribution reveal the basis of metabolic defects observed in shell protein mutants. While a Pdu MCP shell lacking PduA (major BMC hexamer) show similar MCP appearance and distribution to that of WT (Cheng *et al*., 2011, Yang *et al*., 2020, Yang *et al*., 2022), the deletion of EutM (BMC hexamer) of Eut MCP resulted polar body (Fig. 2, Fig. S1). It can be proposed that multiple paralogous BMC shell proteins in Pdu MCP can complement the function of the others to retain the overall shape of the polyhedral structure (Chowdhury *et al*., 2016, Kennedy *et al*., 2021, Yang *et al*., 2020). Unlike Pdu system, the *eut* operon, however, does not synthesize multiple paralogous proteins of same family or the loss perhaps does not get compensated by other shell proteins. It is noteworthy that slower growth phenotype of *eutM* deletion mutant was partially restored by EutK when overexpressed from plasmid (data not shown). The EutK protein possesses single BMC domain like EutM with extra C-terminal extension (Tanaka *et al*., 2010).

While comparing *eutM* with *eutL* mutant, the former might have MCP shells largely fallen apart at the pole together with the enzyme. This resulted large polar body and bright fluorescent signal from exposed EutBC enzyme. It is worth noting that fluorescent signal in the current study demonstrated localization of EutBC enzyme specifically. Unlike *eutM*, a mutation on *eutL* presumably generated comparatively less aberrant shell and produced what is called proto-MCP aggregates that could presumably retain enzyme within it, resulting smaller aggregate and relatively faint fluorescent signal. Similar proto-MCP formation resulting small polar bodies were evident in *pduB* mutant of Pdu MCP (Kennedy *et al*., 2022). It is noteworthy that PduB trimer is structural ortholog to EutL.

It has been observed that the MCPs in *eutN* mutant were all elongated that span between two or more bacterial cells (Fig. 2, Fig. S1), which is closely similar to that in Pdu MCP devoid of PduJ hexamer (Cheng *et al*., 2011, Chowdhury *et al*., 2016). Nevertheless, previous literature revealed some elongated MCP conformation in *pduN* mutants as well, along with enlarged and aggregated phenotypes (Yang *et al*., 2022). The EutN protein was crystalized as hexamer (Tanaka *et al*., 2008). Later, a pentameric conformation of EutN in solution was reported (Wheatley *et al*., 2013). Recent structural polymorphism suggested the possibility of multiple oligomeric state of EutN shell protein (Ochoa *et al*., 2020, Kinney *et al*., 2011).

Another striking feature of Eut MCP is that deletion of *eutK* resulted protein aggregates at both the poles in some cells, a unique phenotype observed under transmission electron microscopy of thin-sectioned sample (Fig. 2). Similar observation was obtained under confocal microscope, where two small fluorescent dots were appeared at both side of the pole of *eutK* mutant cell (Fig. S1). Similar bipolar fluorescent puncta were observed in *mcdA* and *mcdB* mutants of α carboxysomes in *Halothiobacillus neapolitanus*, where defect in spatial distribution was manifested (MacCready *et al*., 2021). Perhaps, EutK also plays similar role in maintaining spatial organization of the Eut MCP. It was greatly noticeable that fluorescent puncta in *eutK* deletion mutant were not localized uniformly in all the cells (Fig. S1). In some cells puncta were at single pole; while in others some puncta in one pole were much brighter than the one in the other pole (Fig. S1). We can surmise that in *eutK* deletion mutant there may be variations in shell and enzyme localization, where shell and enzyme aggregates may either be localized at the opposite poles of the cell or next to each other at the same pole. In some cells, the EutBC enzyme presumably gets aggregated at one pole in the absence of an MCP shell, producing bright fluorescence.

Despite previous studies suggested the importance of EutS shell protein (Choudhary *et al*., 2012, Held *et al*., 2016) in MCP assembly, the deletion of *eutS* does not seem to produce any obvious effect on Eut MCP morphology and distribution.

The growth study (Fig. S2, Fig. 3a) and EA uptake assay (Fig. 4c) indicate that the Eut MCP shell does not appear to impose a permeability barrier to substrates. This is in contrast to the Pdu MCP, where disruption enhances substrate access to the lumen enzyme, resulting faster growth and a growth arrest is manifested when the mutants are subjected to higher B_12_ (Chowdhury *et al*., 2016, Havemann *et al*., 2002, Kennedy *et al*., 2022, Mills *et al*., 2022, Cheng *et al*., 2011, Chowdhury *et al*., 2015, Lehman *et al*., 2017, Yang *et al*., 2022).

Unlike Pdu MCP, the growth defects resulting from Eut MCP shell protein deletion are predominantly due to impairment in the retention of aldehyde intermediate within deformed MCP lumen. Our study revealed substantial rise of acetaldehyde intermediate in the media upon deletion of shell proteins like EutM, -L and -K (Fig. 5, Fig. S3b). The leakage of acetaldehyde from the mutant Eut MCP shell likely reduces carbon availability for downstream enzymes, leading to metabolic deficiencies and growth defects (Fig. 3). This further decreases acetate and ethanol end product formation in the shell protein mutants (Fig. 4a,b). In contrast to *eutM*, -*L* and -*K*, an *eutN* mutant also has growth impairment (Fig. 3a), however, without manifesting any excess acetaldehyde release (Fig. 5). The *eutN* mutant has reduced EA substrate uptake through elongated shell (Fig. 4c), which might be the reason of slower growth phenotype in the said mutant. An *eutS* mutant, on the other hand, did not exhibit severe growth phenotype like other mutants (Fig. 3a), despite leakage of acetaldehyde through MCP shell, resulting lower end-products (acetate and ethanol) (Fig. 4, 5). It can be hypothesized that deletion of EutS produces somewhat porous shell which releases aldehyde, without compromising integrity and morphology of the MCP shell as observed under TEM and confocal microscopic analyses (Fig. 2). EutS is structural ortholog to PduU of Pdu MCP (Tanaka *et al*., 2010). The previous study on Pdu MCP indicated that PduU is a minor shell component, and its deletion exerts only a minimal effect on MCP structure (Havemann & Bobik, 2003, Cheng *et al*., 2011). The relative abundance of EutS was not reported earlier due to absence of Eut MCP purification protocol. However, the near normal morphology of MCPs lacking the EutS shell protein indicates that EutS contributes only minimally to MCP assembly.

We also wanted to assess effect of shell protein mutation on another intermediate, ammonia. The growth studies suggest that the Eut MCP shell of *Salmonella* functions to optimize ethanolamine metabolism exclusively as a carbon source. Within the MCP shell, ethanolamine is converted into acetaldehyde and ammonia (Fig. 1). Here, the amount of ammonia liberated in the media throughout the growth was estimated as stated before (Abdelwahed *et al*., 2022). It is evident that the mutant shells liberate almost similar amount of ammonia to that of WT (Fig. S4), indicating indispensable role of Eut MCP shell while serving EA as nitrogen source.

It is pertinent to mention that Eut shell protein mutants were not fully complemented by ectopic expression from plasmid. The growth of *eutM*, -*N* and -*L* shell protein mutants could only be partially complemented even after addition of as high as 500 µM IPTG, except in case of *eutK* (Fig. S5). The growth of *eutK* mutant was recovered fully at as low as 20 µM IPTG. Due to small phenotype the growth complementation study was not performed in case of *eutS* mutant (Fig. S5). Strategy of complementation among major shell proteins often fails due to improper dosage of the protein expressed from plasmid (Chowdhury *et al*., 2016, Yang *et al*., 2025, Yang *et al*., 2022). Additionally, due to protein–protein interactions, the deletion of a single protein could potentially lead to the loss of multiple associated proteins including luminal enzymes (Held *et al*., 2016, Jallet *et al*., 2024, Jorda *et al*., 2015, Yang *et al*., 2025), which could also explain the failure of complementation by a single protein. Recently, successful restoration of bacterial physiology has been achieved through ectopic expression of the shell protein in multiple Eut shell protein mutants, which validate the generation of non-polar genetic mutants (Bhalerao *et al*., 2025). Also, gene expression study (Fig. S5b) with semi quantitative RT PCR analyses has ruled out plausible polarity in the mutant preparation.

Further, crude enzyme activity analyses thoroughly deliver holistic insights on shell and enzyme assembly. It was evident that enzyme encapsulation was significantly affected by the loss of individual shell proteins. However, impact was more profound in *eutM* and *eutL* mutants, indicating pivotal role of EutM hexamer and EutL trimer in maintaining shell integrity and cargo loading. The consequence of *eutM* and *eutL* deletion on bacterial physiology has been elucidated recently (Bhalerao *et al*., 2025). In comparison to *eutM*, the effect of *eutL* was less pronounced, likely due to its less abundance as compared to the former. Some enzyme cargo like EutBC might still be attached within proto-MCP aggregate of *eutL* mutant resulting similar ethanol to WT (20% reduction, Fig. 4b). But markedly reduced acetaldehyde dehydrogenase (EutE) activity indicates poor enzyme retention, thereby diminishing acetate production in this mutant (Fig. 4a, 6). The deletion of *eutK*, on the other hand results very low EutBC activity indicating poor retention of the enzyme. However, the deletion of this shell protein exerts only a minimal impact on EutE activity (Fig 6). We therefore surmise that the reduced acetate formation in the *eutK* mutant is likely resulting from poor retention of the acetaldehyde intermediate within the microcompartment shell. In contrast, enzyme activities in the *eutS* deletion mutant are comparable to those in the wild type, indicating that enzymes are properly retained within the MCP lumen.

Interestingly, the *in vitro* enzyme activities were diminished in *eutN* mutant; however, the mutant is able to form similar acetate and ethanol end product to WT. The function of EutN is to close geometric curvature of facets and seal vertices to introduce roughly icosahedral geometry of the MCP shell. A deletion of *eutN* thus induce an elongated flat sheet. The protein sheet has been able to serve as a scaffold framework to a number of enzymes (Zhang *et al*., 2019). The confocal microscopic analyses of *eutN* mutant indicate EutBC enzyme extends across elongated sheet. We speculate that the enzymes scaffolded onto protein sheet in *eutN* mutant function properly (Fig. 4). The reduced *in vitro* enzyme activity in *eutN* mutant might presumably be due to collapse of protein sheets followed by dilution of enzymes during cell lysis.

Collectively, these findings provide insights into the differential roles of shell proteins in EA metabolism within Eut MCP. Deciphering functional role of Eut MCP shell proteins would open up paradigms that underlie MCP functional diversification and advance fundamental understanding of these unusual and important bacterial organelles in microbial ecology and metabolism.

## Experimental procedures

### Chemicals and reagents

Ethanolamine hydrochloride, Cyanocobalamin (Vit. B_12_), glycerol, 3-Methyl-2-benzothiazolinone hydrazone hydrochloride (MBTH), were from Sigma-Aldrich (St. Louis, MO). Lysogeny broth, antibiotics, Isopropyl-β-D-1-thiogalactopyranoside (IPTG), and arabinose were from Himedia (Mumbai, India). Q5 DNA polymerase, restriction enzymes, and T4 ligase were from New England Biolabs (Beverly, MA). 2X PCR TaqMixture^TM^ was from Himedia (Mumbai, India). Other chemicals were obtained from Qualigens Thermofisher Scientific (Powai, India).

### Bacterial strains, media, and growth conditions

The bacterial strains used are listed in Table S1. All strains are derived from *Salmonella enterica* serovar Typhimurium strain LT2. The rich medium used was Lennox (lysogeny broth) (LB) medium (Himedia, Mumbai, India). For selecting transformants TYE media was used; where bacto-tryptone, yeast extract, and agar were purchased from Himedia Laboratories (Himedia, Mumbai, India). The minimal media used were no-carbon-E (NCE) and no carbon-no nitrogen (NCN) medium (Bhalerao *et al*., 2025). Minimal media were supplemented with 1 mM MgSO4, 0.3% each of valine, isoleucine, leucine and threonine and 50 µM ferric citrate. CN-B_12_ or Vit. B_12_ (20 and 150 nM) was served as exogenous B_12_ source. Antibiotics were used at the following concentrations: ampicillin, 100 μg ml^-1^ and chloramphenicol, 20 μg ml^-1^. Solid media were prepared by adding agar (1.5%, Himedia) to LB, TYE, NCE, or NCN medium. All the growth studies were performed in aerobic conditions.

### Construction of deletion and insertion mutants and complemented strains

All primers and plasmids used in this study are listed in Table S2. Chromosomal deletion of individual Eut MCP shell proteins was constructed by recombineering as stated earlier (Chowdhury *et al*., 2016, Datsenko & Wanner, 2000). For clean mutant preparation the mutant gene was transduced to the cell with no mutation background by P22 HT105-int mediated transduction. For double mutant preparation the Cam cassette at the respective gene loci was transduced to another mutant. Transductants were tested for phage contamination and sensitivity by streaking on green plates against P22 H5 (Cheng *et al*., 2011). For complementation studies, the genes encoding Eut shell proteins were cloned into the pLac22 plasmid within the *Bgl*II and *Hin*dIII sites.

To insert m-Cherry cassette at the C-terminus chromosomal locus of *eutC* gene (R298), a dual-selectable mpheS-kan cassette was first amplified with 40 bp target-homology flanks. The linear PCR amplicon was introduced by electroporation into the λ-RED expressing host strain. Recombinants were selected on kanamycin plates and verified for correct genomic integration via colony PCR. A PCR-amplified m-Cherry gene containing upstream and downstream homology flank was electroporated into the cassette-containing strain. Transformants were counter-selected on media containing DL-4-chlorophenylalanine. Resistant colonies were screened for kanamycin sensitivity to confirm the loss of the mpheS-kan cassette. Final mutant constructs were confirmed by PCR amplification and DNA sequencing.

### Microscopic analyses of Eut MCP morphology

Overnight culture of wild-type and shell protein mutants were diluted in 1:100 ratio in minimal media supplemented with 30 mM ethanolamine and 150 nM CN B_12_ to express Eut microcompartment. The culture was allowed to grow further until mid log phase. After harvesting, the bacterial cells were allowed to fix with 2.5% glutaraldehyde and 2% paraformaldehyde mixture prepared in 0.1 M cacodylate buffer for 2 h. This was followed by staining with 1% osmium tetroxide in a same buffer for 2 h. En-bloc staining was carried out with 2% uranyl acetate for 1 h, followed by subsequent dehydration steps through a graded ethanol series. Samples were embedded in Araldite-502, dodecenylsuccinic anhydride and epoxy accelerator DMP-30 and polymerized at 60°C for 72 h. After embedding, the thin sections of ∼70 nm thickness were made by ultramicrotome (Leica UC7). The sections were mounted on the copper grid and stained with 0.1% (w/v) Reynolds lead citrate for 8 min. The stained sections were subjected to visualization under transmission electron microscope (JEM 1400 PLUS, JEOL, Japan).

To better visualize Eut MCP through confocal microscope, an m-Cherry reporter gene was fused at the 3’-end of *eutBC* encoding ethanolamine ammonia lyase, the first enzyme of ethanolamine metabolism by mPheS-Kan mediated recombineering method as stated above. The bacteria were subjected to express Eut MCP by growing in NCE media, supplemented with ethanolamine and B_12_ as stated before.

### Growth studies

Growth studies were performed at both limiting (20 nM) and saturating (150 nM) CN-B_12_ as previously described (Chowdhury *et al*., 2015, Chowdhury *et al*., 2016) with a BioTek Epoch 2 Microplate Spectrophotometer (Agilent-BioTek, USA). Doubling times were calculated from semilog plots (Cheng *et al*., 2011). For determination of levels of metabolites produced, 50 ml cultures were grown in NCE media supplemented with 30 mM EA and 150 nM CNB_12_ in 250 ml Erlenmeyer flasks at 37°C with shaking at 250 rpm. These cultures were inoculated to an optical density at 600 nm (OD_600_) 0.1 with an LB overnight culture that had been centrifuged and resuspended in NCE with 1 mM MgSO4. The growth study in the plate reader was performed with six replicates and three biological replicates.

### Quantification of substrate uptake and product release

Cultures of wild-type and shell protein mutants were sampled at timed intervals of 3 h. Cells were removed by centrifugation followed by filtration with 0.22 μm syringe filters. Acetate and ethanol, the end products of EA metabolism released in the culture media were estimated by high-performance liquid chromatography (HPLC) using a Bio-Rad Aminex HPX-87H (300 × 7.8 mm) column eluted isocratically with 5 mM H_2_SO_4_ as described elsewhere (Chowdhury & Bobik, 2019). Xylitol (10 mM) was used as internal standard. The elution was monitored through RI detector (Hitachi LaChrom Elite HPLC). The uptake of ethanolamine substrate was estimated after derivatizing with dansyl chloride (Thiennimitr *et al*., 2011), followed by running through reverse phase UPLC (Shimadzu) using a Waters C18 XBridge BEH C18 Column (130Å, 3.5 µm, 4.6 mm × 150 mm) and eluted isocratically with acetonitrile: water (50:50, v/v) at a flow rate of 0.6 ml min^-1^. The compound was detected through fluorescence (ex: 375 nm, em: 525 nm) (Chen *et al*., 1981, Mir-Cerda *et al*., 2022).

### Estimation of acetaldehyde intermediate

Acetaldehyde is one of the key intermediates, which must be retained within microcompartment (MCP) to keep EA metabolism up and running. The lesser the efficiency of the mutant shell to retain the acetaldehyde, the more will be its released in media, which can be estimated spectrophotometrically at 305 nm after reacting with MBTH (3-Methyl-2-Benzothiazolinone Hydrazone).

The acetaldehyde released from WT and Eut shell protein mutants was estimated in both shake and crimp-sealed containers. In brief, the cultures of WT and mutants were grown in 250 ml Erlenmeyer flasks for 36 h. The sample (50 µl) was aliquoted at timed intervals of 3 h. The amount of acetaldehyde remaining was estimated by reacting with 3-Methyl-2-Benzothiazolinone Hydrazone (MBTH) and measured at 305 nm as described previously (Ochoa *et al*., 2023, Penrod & Roth, 2006). In case of sealed container, the 10 ml culture was incubated in 100 ml sterile glass vials containing rubber caps were sealed with aluminum caps with the help of crimper tool. The glass vials were incubated at 37°C for 36 h. The amount of acetaldehyde released out the shell was estimated as stated before.

### Enzyme activity

The strains to be analyzed were primarily grown in Luria broth for 16 h. Secondary inoculation was carried out in minimal media containing No carbon essential (NCE) media supplemented with 30 mM ethanolamine, 150 nM vitamin B_12_ and 20 mM glycerol. Appropriate controls were also set up and these cultures were grown at 37 for 7 h under shaking conditions at 250 rpm.

Cell lysis was carried out as described previously with minor modifications (Trettel & Winkler, 2023). The cell pellets were first resuspended in cell lysis buffer (50 mM Tris HCl, 1 mM KCl, pH 8.0) supplemented with BPER II (Thermo Scientific). The suspension was incubated at 37 °C for 1 h, followed by brief sonication for 2 min with alternating 10 s pulses and 10 s intervals. The whole cell lysate was used for crude enzyme assay analysis.

Coenzyme B_12_-dependent ethanolamine ammonia lyase (EutBC)) activities from crude cell lysate was determined by a linked continuous spectrophotometric assay, as described previously (Akita *et al*., 2010). The EutE acetaldehyde dehydrogenase activity was measured by following the conversion of NAD^+^ to NADH spectrophotometrically at 340 nm, as described previously (Leal *et al*., 2003, Lehman *et al*., 2017).

## Acknowledgements

This study was supported mainly by the ad-hoc grant from the Indian Council of Medical Research (ICMR), Govt. of India [Diarr/Adhoc/5/2022-ECD-II] to C.C. M.R.B. and A.A.S acknowledge the Council of Scientific and Industrial Research (CSIR) and Mahatma Jyotiba Phule Research Fellowship (MJPRF) for fellowships. A.S.D. and Y.D.S thank for their fellowship support from Department of Biotechnology (DBT), Govt of India, and ICMR, Govt. of India, respectively. Authors acknowledge The Advanced Centre for Treatment, Research and Education in Cancer (ACTREC), Tata Memorial Centre (TMC), Navi Mumbai for transmission electron microscopic analyses.

## Supporting information

Additional supporting information may be found in the online version of this article at the publisher’s web-site

## Data Availability Statement

Any additional information required to reanalyze the data reported in this study can be made available upon request.

