## Supporting information for "Genetic analyses reveal distinct functional roles of Eut microcompartment shell proteins in ethanolamine metabolism in *Salmonella*"

**Tables**

**Table S1**. Bacterial Strains used in the study

| **Species/Strain** | **Genotype** | **Source** |
| --- | --- | --- |
| ***E. coli*** | | |
| **YS9** | DH5alpha/pKD46 | Lab Collection |
| **YS10** | DH5alpha/pCP20 |  |
| **MB1** | DH5alpha/pLac22 |  |
| **CE28** | *E. coli*/ pKD4 |  |
| **CE29** | *E. coli* BW25141/pKD3 |  |
| **CE30** | *E. coli* top10/pJD141 (Template for mPheS-kan) |  |
| **CE3** | WT/pLac22 |  |
| **CE1** | *Salmonella enterica* serovar Typhimurium LT2 |  |
| **CE35** | ∆*eutK*::frt |  |
| **CE34** | ∆*eutL*::frt |  |
| **CE37** | ∆*eutM*::frt |  |
| **CE36** | ∆*eutS*::frt |  |
| ***Salmonella enterica* Typhimurium LT2** | | |
| **RD8** | ∆*eutB*::frt | This Study |
| **MB7** | ∆*eutN*::frt |  |
| **AD24** | Δ*eutK*::frt/pLac22 |  |
| **AD8** | ∆*eutK*::frt/pLac22-*eutK* |  |
| **MB34** | Δ*eutL*::frt/pLac22 |  |
| **MB33** | ∆*eutL*::frt/pLac22-*eutL* |  |
| **AD25** | Δ*eutM*::frt/pLac22 |  |
| **AD26** | Δ*eutM*::frt/pLac22-*eutM* |  |
| **MB36** | Δ*eutN*::frt/pLac22 |  |
| **MB30** | Δ*eutN*/pLac22-*eutN* |  |
| **MB38** | Δ*eutS*::frt/pLac22 |  |
| **MB37** | Δ*eutS*/pLac22-*eutS* |  |
| **AD43** | ∆*eutK*::frt-∆*eutM*::frt-Cam-frt |  |
| **AD39** | DH5a/pLac22-*mCherry* |  |
| **AD53** | *eutC*-R298::*mCherry* |  |
| **AD89** | *ΔeutK::*frt*-eutC-*R298:*:mCherry* |  |
| **AD175** | Δ*eutL*::frt- *eutC-*R298*::mCherry* |  |
| **AD93** | *ΔeutM:*:frt- *eutC-*R298*::mCherry* |  |
| **AD173** | Δ*eutN*::frt- *eutC-*R298*::mCherry* |  |
| **AD133** | *ΔeutS:*:frt- *eutC-*R298*::mCherry* |  |
| **AS03** | Δ*adhP*::frt |  |
| **AS10** | Δ*eutB*::frt-Δ*adhP*::frt-cam-frt |  |
| **AS05** | Δ*eutM*::frt-Δ*eutB*::frt-cam-frt |  |
| **AS17** | Δ*eutN*::frt-Δ*eutB*::frt-cam-frt |  |
| **AS20** | Δ*eutS*::frt-Δ*eutB*::frt-cam-frt |  |
| **AS23** | Δ*eutL*::frt-Δ*eutB*::frt-cam-frt |  |
| **AS26** | Δ*eutB*::frt-Δ*eutK*::frt-cam-frt |  |

**Table S2**. Primers used in this study

| **Description** | **Primer** | **Sequence (5’-3’)** |
| --- | --- | --- |
| **Mutant Preparation** | | |
| ***eutK*** | *eutK-Cam-F* | TGCTGGATATCGCCCGTAACCCTGTCCAGCGTGCGTAACGGAGGCTGCCATGTAGGCTGGAGCTGCTTCG |
|  | *eutK-Cam-R* | CATGATCTTCTATGCCGGACGCCCGCCAGGACGCCCGGCCCTCCGACAGAATATGAATATCCTCCTTAGTTC |
| ***eutL*** | *eutL-Cam-F* | TGCTGGAGCAGAAAGCGTCCGGCATCAACATGACCCGTTAAGGAGACATCTGTAGGCTGGAGCTGCTTCG |
|  | *eutL-Cam-R* | ATTCCGTCCACTTCCAGTAATCCCAGGGCATTGATCATTGGCAGCCTCCGATATGAATATCCTCCTTAGTTC |
| ***eutM*** | *eutM-Cam-F* | GGTTGAATGAACGGTCCCGTTCTGGACCCCTTTAATGAGAGGAAAACACGTGTAGGCTGGAGCTGCTTCG |
|  | *eutM-Cam-R* | GCGCGCATTCATGCCGGATGGCGACGCTGCGCGTCTTATCCGGCCTACCAATATGAATATCCTCCTTAGTTC |
| ***eutN*** | *eutN-Cam-F* | CATTGCCTGATGGTGCGCAGATTCAAGGCCTACGACCCAACGGGCAATCCTGTAGGCTGGAGCTGCTTCG |
|  | *eutN-Cam-R* | TACCGCTTTCACCACCTGTTCAATATCCTGTTGATTCATGATGTTCAATCATATGAATATCCTCCTTAGTTC |
| ***eutS*** | *eutS-Cam-F* | CCGACAAAAAATTGCCACGATGACGGCAGTTTCAGTGGAGACGGTGAGCATGTAGGCTGGAGCTGCTTCG |
|  | *eutS-Cam-R* | CCGGCACCGACCGCGCCGACAAAAGCAATACGTTTCATTACCGCCACCAAATATGAATATCCTCCTTAGTTC |
| ***eutB*** | *eutB-Cam-F* | GGTGAAATCACTCGCATTTCCTTCCTGAGGGAACGACTTTGTAGGCTGGAGCTGCTTCG |
|  | *eutB-Cam-R* | TCAATCTGTTTTTGATCCATGGTGTTATCCCCGCGTCAATATGAATATCCTCCTTAGTTC |
| ***eutC*-R298::mPhes** | *eutC*-R-298-mPhes-F | TGGCCAAACGGATGCTGGAGCAGAAAGCGTCCGGCATCAACATGACCCGTGAATTCGCGGCCGCATCTAG |
|  | *eutC*-R-298-mPhes-R | GCAGTCACTGAAGGTCGAATTAAATCTAATGCAGGCATGATGTCTCCTTACTGCAGCGGCCGCTACTAGT |
| ***eutC*-R298::mCherry** | *eutC C-ter mCherry-F* | GGAGCAGAAAGCGTCCGGCATCAACATGACCCGTGGCTCTATGGTGAGCAAGGGCGAGG |
|  | *eutC C-ter mCherry-R* | TCGAATTAAATCTAATGCAGGCATTGGCTAACCTCCTTTATTACTTGTACAGCTCGTCC |
| ***adhP*** | *adhP*_Cam_F | CTATGCAGTTCACTATTAATAAAAGGAGAAGGTT TGTAGGCTGGAGCTGCTTCG |
|  | *adhP*_Cam_R | TCTGCTGTCTGTATACTCAGACGACCACAGCAAATCGT |
| Mutant and clone confirmation | | |
| **pLac** | PLac-F | CCCCAGGCTTTACACTTT |
|  | PLac-R | GTTAGATTTCATACACGGTGCCTG |
| ***eutK*** | *eutK*-F | TCGTACCTCTCGTCTACCGCA |
|  | *eutK*-R | TCATCAAGCAGGATTTCCGT |
| ***eutL*** | *eutL*-F | GGCAGTCGGAAAGCCTTT |
|  | *eutL*-R | CGACTAATCACACGGCCTGT |
| ***eutM*** | *eutM*-F | AAATCACTCAGCGTCTGGGA |
|  | *eutM*-R | TCATTTCCACCATCAGCAAT |
| ***eutN*** | *eutN*-F | TGCAAAGCCGCAACCGAT |
|  | *eutN*-R | TTTCGCTGACGGCAAGTT |
| ***eutS*** | *eutS*-F | CGGAACAGGCTTAAATTCAGA |
|  | *eutS*-R | TGCAACGTGGTAATTAAGGC |
| ***eutB*** | *eutB -F* | TACGCCTCTTTTTGGCGGATCGGTT |
|  | *eutB -R* | CTGTCCCATTGACGCCATCACGCTA |
| ***adhP*** | *adhP -F* | TACGTACCCGGCGGTGAATG |
|  | *adhP -R* | AGAGCGAAGAAGGTATTCAGTAATTCTT |

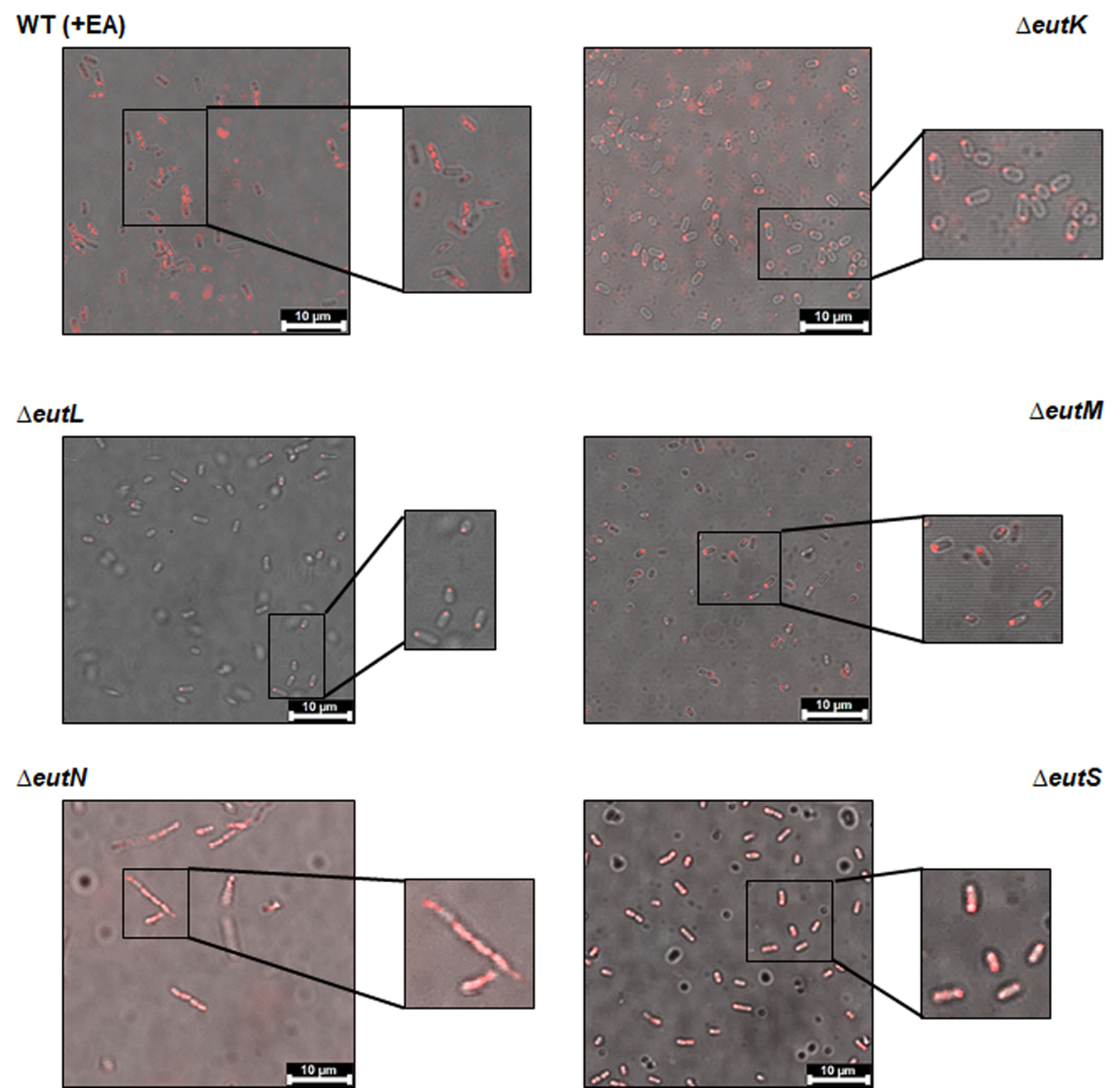

Figure S1. In-situ localization of Eut MCP in wild type *Salmonella* and Eut shell protein mutants grown in presence of EA and B_12_. Scale bar is 10 µm.

Figure S2. Growth of wild type and Eut shell protein mutants in no-carbon E minimal medium supplemented with EA and limiting B_12_ concentration (20 nM). The error estimate shown is ± one standard deviation and are based on three biological replicates. A one-way ANOVA was used for statistical significance. **p < 0.05.

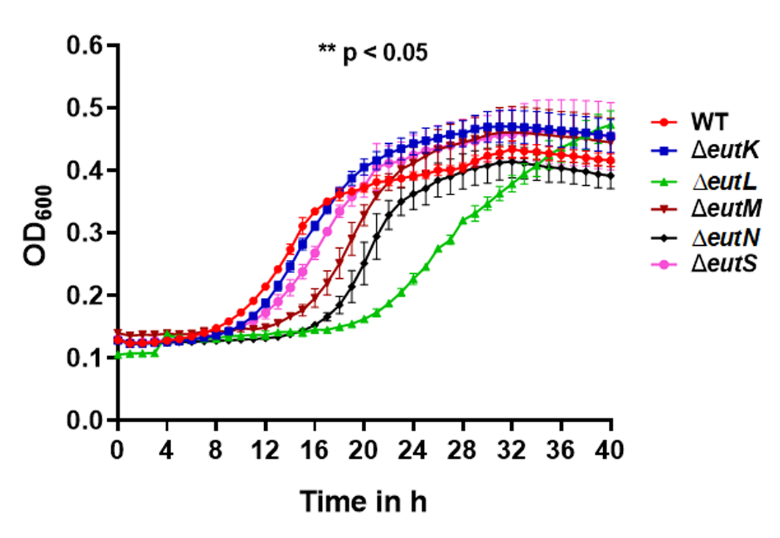

a)
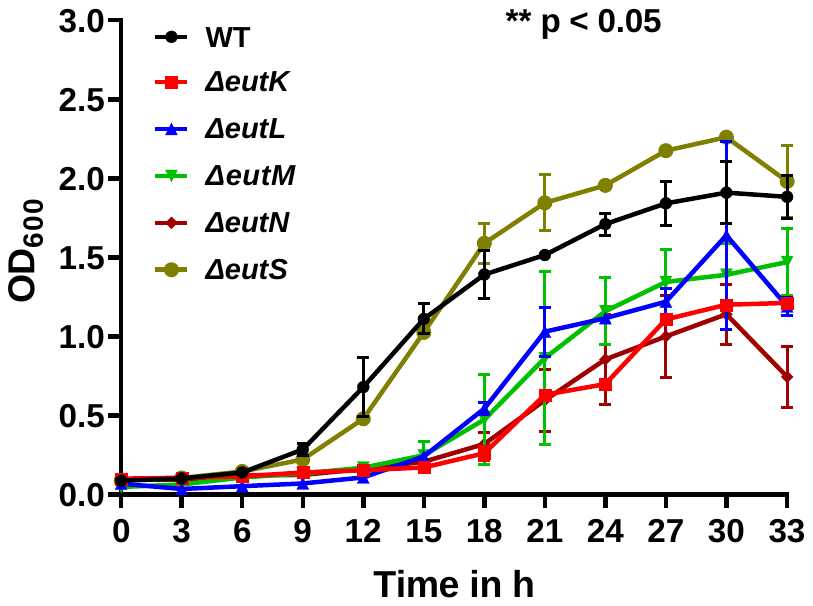

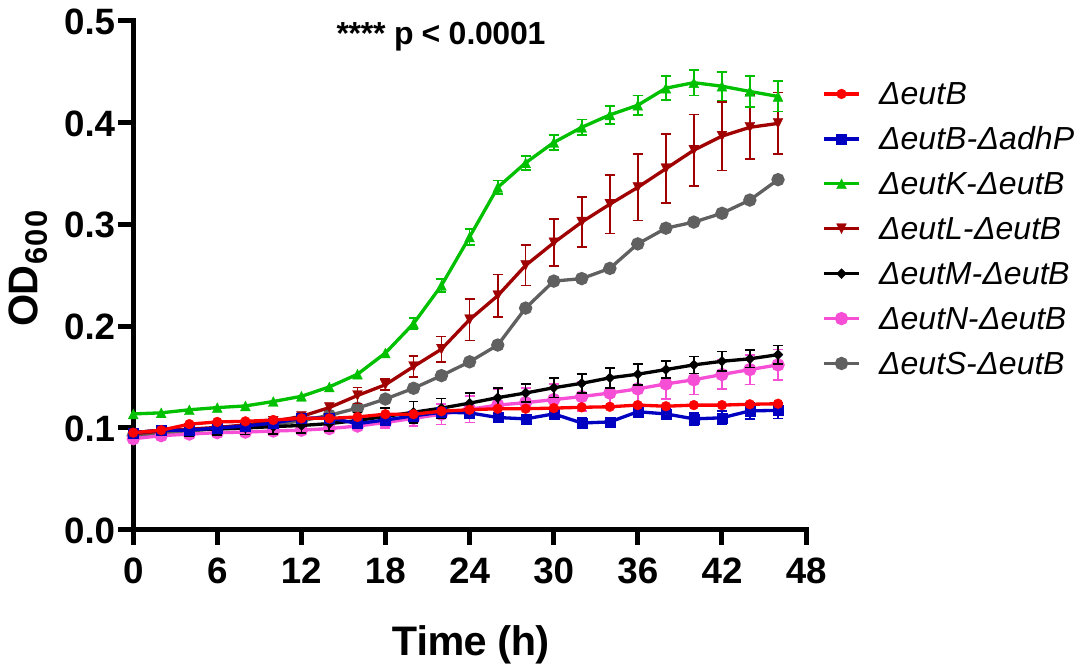
 b)

Figure S3. a) Growth of WT and shell protein mutants in NCE minimal media supplemented with 30 mM EA (as carbon source) and 150 nM CNB_12_ in 250 ml Erlenmeyer flasks at 37°C with shaking at 250 rpm. b) Growth of wild type and Eut shell protein mutants on ethanol (20 mM). The shell protein mutation breaks the shell of the Eut MCP allowing acetaldehyde to reach MCP lumen enzymes by free diffusion. The *adhP* lacks the cytoplasmic AdhP alcohol dehydrogenase which converts ethanol to acetaldehyde. An *eutB* mutant background has been taken to block utilization of ethanolamine. However, this will not block the expression of Eut MCP. The error estimate shown is ± one standard deviation and are based on three biological replicates. A one-way ANOVA was used for statistical significance. **p < 0.05, ****p < 0.0001

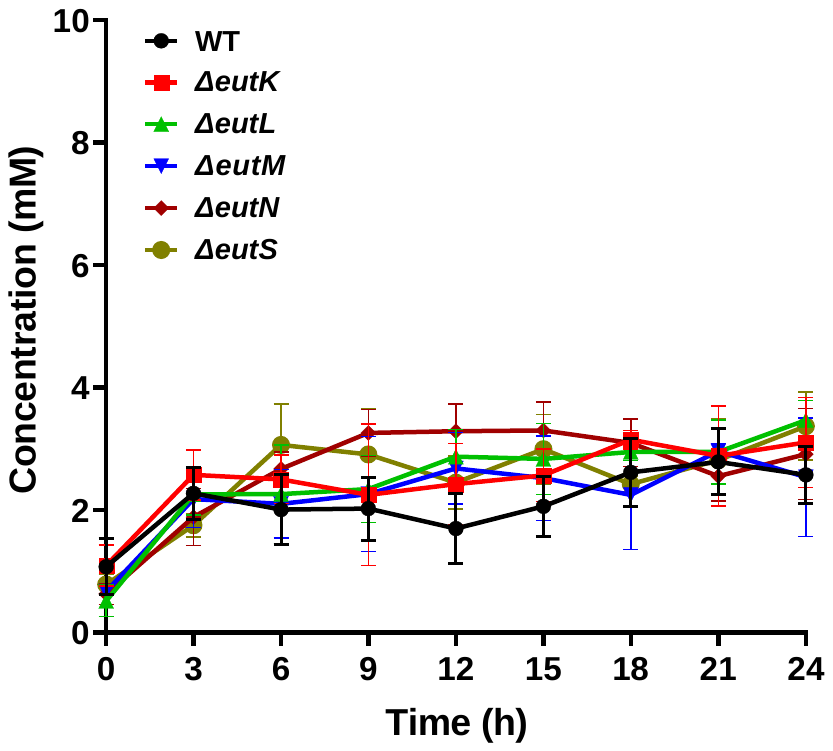

Figure S4. Estimation of ammonia liberated into culture media. All the shell protein mutants excrete similar amount of ammonia to WT. The error estimate shown is ± one standard deviation and are based on four biological replicates. A one-way ANOVA was used for statistical significance. NS p > 0.1

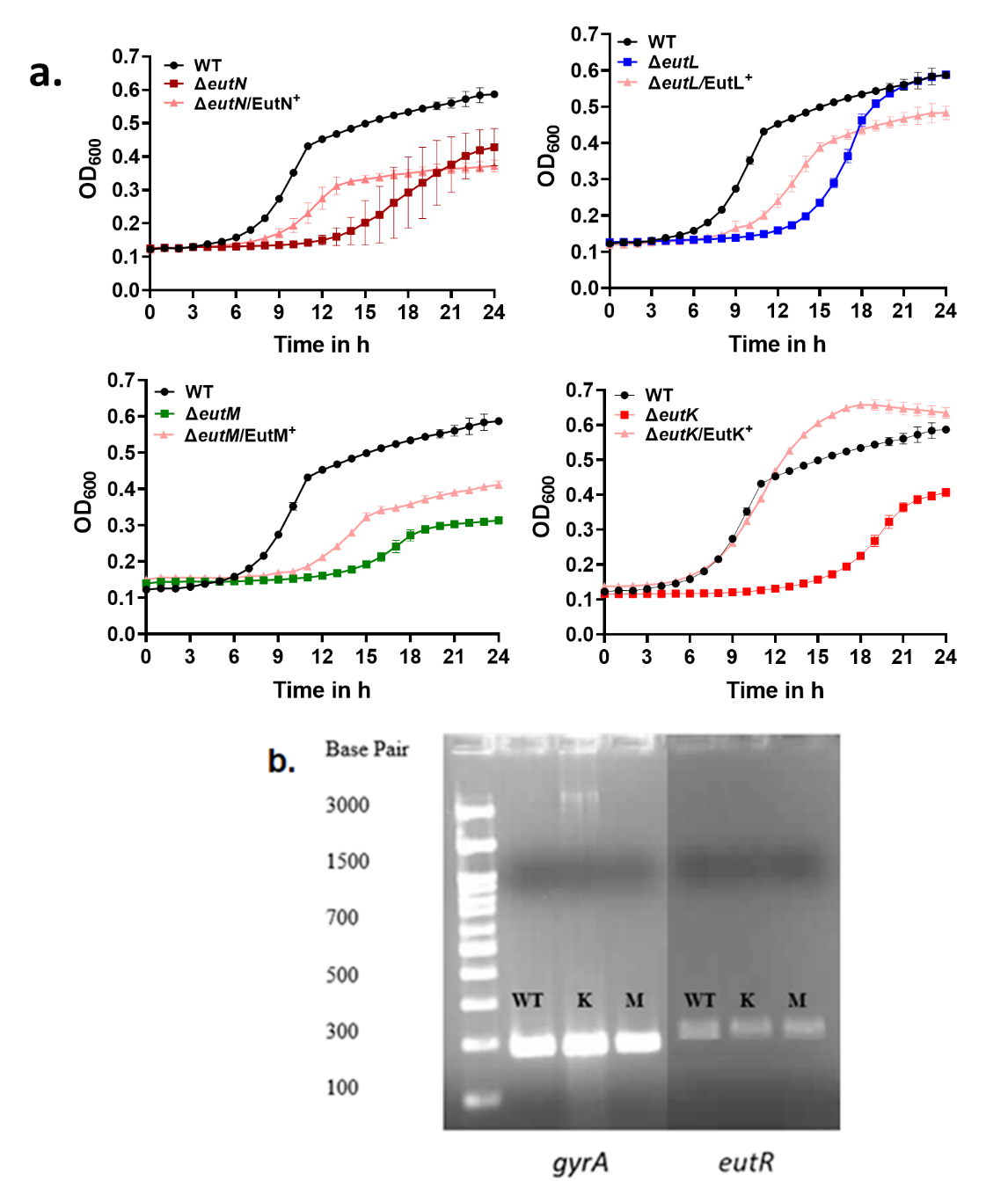

Figure S5. a) Complementation of growth of shell protein mutants by corresponding genes expressed from plasmid. b) The expression of *eutR* in the *ΔeutK* and *ΔeutM* is similar to that of the wild type indicating that these mutants are non-polar. *gyrA* has been used as house-keeping gene control.
